# 7 Tesla fMRI characterisation of the cortical-depth-dependent BOLD response in early human development

**DOI:** 10.1101/2025.08.18.670552

**Authors:** Jucha Willers Moore, Philippa Bridgen, Elisabeth Pickles, Pierluigi Di Cio, Lucy Billimoria, Ines Tomazinho, Cidalia Da Costa, Dario Gallo, Grant Hartung, Alena Uus, Maria Deprez, Sharon L. Giles, A. David Edwards, Jo V. Hajnal, Shaihan J. Malik, Jonathan R. Polimeni, Tomoki Arichi

## Abstract

Human cortical development leading up to and around birth is crucial for lifelong brain function. Cortical activity can be studied using BOLD fMRI, however, previously limited sensitivity and spatial specificity has constrained understanding of how its emergence relates to functional cortical circuitry and neurovascular development at the mesoscale. To resolve this, we used ultra-high-field 7 Tesla MRI to acquire sub-millimetre resolution BOLD-fMRI data from 40 newborns and 4 adults. In all subjects, passive right-hand movement elicited localised, positive BOLD responses in contralateral primary somatosensory cortex. In newborns, depth-specific BOLD responses were still evident in the thinner cortex, with developmental changes in response temporal features and amplitudes at different depths. This provides insight into key rapidly evolving factors in early cortical development including neuronal function, vascular architecture, and neurovascular coupling. Our framework and findings provide a foundation for future studies of emerging cortical circuitry and how disruption leads to adverse outcomes.

## Introduction

Development of the cortex and its intricate framework of neuronal circuitry underling complex processing across the lifespan, begins early in gestation and is maximal in the perinatal period. Functional MRI (fMRI) is increasingly used to provide non-invasive insight into emerging patterns of activity in the developing human cortex. Studies in preterm infants have demonstrated rapidly evolving patterns of sensory activation using fMRI, moving from predominately the primary sensory cortex only to increasingly complex and hierarchical patterns with advancing post-menstrual age (PMA)^1^. However, fundamental insight into how neuronal activity, cortical circuits and haemodynamic responses are established during this critical period are still lacking. This is largely due to the low spatial resolution in typical fMRI studies acquired at 3 Tesla (T) with standard voxel volumes of 8–27 mm^3^, preventing characterisation of fine-scale cortical processing across its emerging layers. Interpretation of fMRI studies in infants are further confounded by concomitant development of vascular architecture, cerebral physiology and neurovascular coupling, all of which are rapidly changing and likely significantly impact on the blood-oxygenation-level-dependent (BOLD) fMRI response acquired^2^.

The perinatal development of human cortical anatomy and structural connectivity have been characterised using postmortem histological assessments and non-invasively with neuroimaging. However, much less is known about perinatal functional development and its relationship with concomitant structural maturation, thus functional neuroimaging at spatial scales relevant to these rapid changes in brain organization is needed. Early in the third trimester of gestation, the cerebral cortex’s six canonical layers are broadly identifiable, with thalamocortical afferents entering the cortical plate and forming connections with the future Layer IV neurons from ∼26 gestational weeks (GW)^3^. After 36 GW, callosal and long-range association fibres reach the cortex and by the end of gestation, all major white-matter tracts are in place^4^. Around this time, extensive growth of intracortical connections and short-range corticocortical fibres begins^5^, essential for brain-wide integration and coordination of information processing. This widespread development makes the neonatal cortex particularly vulnerable to disruption and injury, as evidenced by the detrimental effects of prematurity including cortical structural abnormalities^6^ and later adverse cognitive outcomes^7,8^. fMRI may thus improve mechanistic understanding of the relationship between altered early cortical development, brain function and later neurodevelopment.

Although fMRI can provide valuable insight into the emergence of cortical function, it is known that key factors that influence BOLD-based measures of brain activity—namely, the vasculature, cerebral physiology and neurovascular coupling—are also undergoing rapid and considerable development during the neonatal period^2^. As a result, the neonatal BOLD response differs markedly from adults, with a slower time to peak, broader temporal dispersion, and decreased amplitude^9^. Furthermore, interpretation of BOLD signal acquired at conventional field strengths is likely influenced by low acquisition resolutions, resulting in inclusion of non-specific BOLD signal from large draining veins^10^. fMRI acquired at ultra-high field (≥7 T) markedly increases signal-to-noise and functional-contrast-to-noise ratios (SNR and fCNR)^11^, enabling dramatically higher spatial resolution (submillimetre voxel sizes) than conventional MR systems. Additionally, higher field strengths increase the relative contribution to BOLD signal from the microvasculature^12^. BOLD responses are therefore better localised to cortical grey matter at 7 T compared to those measured at 3 T^11^. This high spatial resolution enables functional activity to be resolved across cortical depths, columns, and small subcortical structures^13,14^. Furthermore, voxel vascular compartment compositions are more heterogenous, with some containing large pial veins and others not. Thus spatiotemporal differences of the BOLD response across cortical depths can inform about different levels of the intracortical vascular hierarchy^15^ and neuronal circuitry.

Although 7 T MR neuroimaging is becoming increasingly common in adult studies, neonates present additional practical and MRI sequence design challenges^16,17^. These include accounting for increased specific absorption rate (SAR) and temperature instability^18^, as well as adapting sequence parameter values for the marked differences in relaxation rates across age^19^. Furthermore, reduced cortical thickness (2 mm in neonates vs. 3 mm in adults)^20^ presents an even greater challenge for discriminating fine-scale cortical activity. However, the potential benefits still far outweigh the challenges, as the increased spatial resolution of 7 T fMRI may provide fundamental new insights into how neuronal activity, cortical circuits and haemodynamics are established during the critical neonatal period.

Motivated by this, we developed a platform for high-resolution 7 T BOLD fMRI to study haemodynamic responses in 40 neonates aged 33.57–46.43 weeks PMA and 4 adults during passive sensorimotor stimulation. BOLD responses were characterized in the primary somatosensory cortex (S1) across three cortical depths representing different levels of the vascular hierarchy and cortical circuitry. Given known differences in vascular anatomy, physiology and neuronal activity in development, we hypothesised that neonatal BOLD response profiles would differ from those in adults. Based on previously characterised spatiotemporal developmental gradients of neuronal and vascular architecture across cortical depths, we further hypothesised that cortical-depth-dependent BOLD response profiles would change across development.

## Results

### Sensorimotor stimulation produces distinct BOLD responses in neonates and adults using 7 T fMRI

fMRI data were collected from 40 infants aged 33.57–46.43 weeks PMA (median: 37.71 weeks PMA, 22 female; Table 1) recruited from the postnatal and neonatal wards at St. Thomas Hospital London and 4 adults aged 22–27 years old (median: 25.5 years, 3 female) on a 7 T MRI system with a 1Tx-32Rx head coil in neonates and an 8Tx-32Rx head coil in adults. Gradient-Echo (GRE)-BOLD fMRI data were acquired with 0.8-mm isotropic resolution and a restricted field-of-view containing S1. Functional responses were induced by 26.6-s blocks of sensorimotor stimulation comprising right wrist passive extension and flexion using an MR compatible pneumatic robotic device (Figure 1). Neonates were scanned in natural sleep. Overall, 7 T scanning was tolerated well; neonates remained clinically stable with no adverse events during data collection.

**Figure 1.**
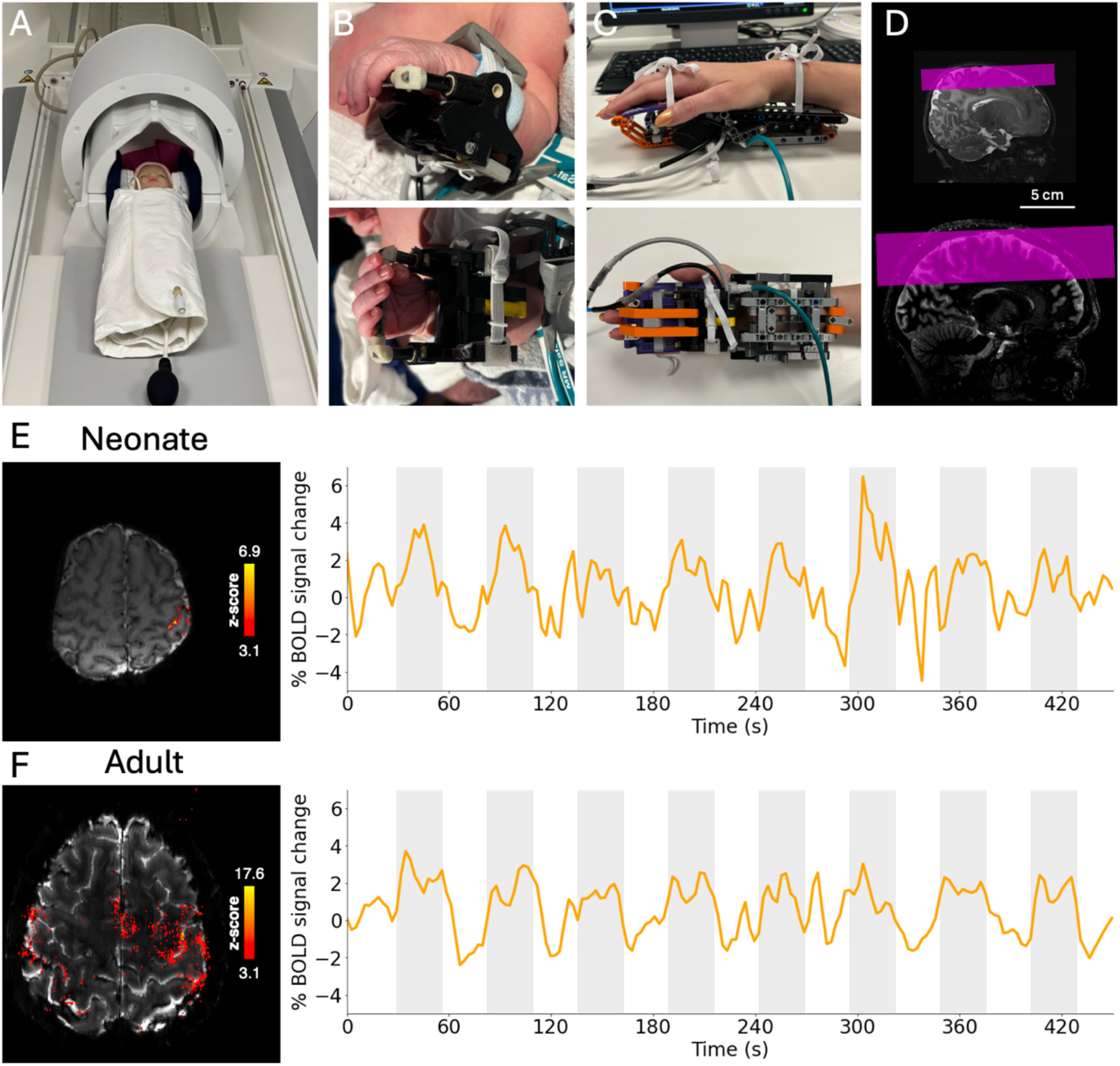
Setup for fMRI data acquisition in neonates on the 7 T MRI system. (A) Demonstration of neonate positioning inside the head coil at isocentre by foam cushions placed around the head. Infants are immobilised in a vacuum-evacuated blanket, and inflatable cushions were placed either side of the infant’s head to further prevent head motion. The MR-compatible pneumatically-driven wrist device which provides passive stimulation of the wrist inside the MRI scanner, fitted onto (B) a neonate wrist (C) and an adult wrist. (D) Exemplar fMRI acquisition field-of-view (pink) in neonates (top) and adults (bottom) overlaid onto a T_2_-weighted anatomical image. (E, F) Passive movement of the right wrist produces localised activation in the contralateral (left) primary somatosensory cortex in (E) one exemplar neonate (38 weeks PMA) and (F) adult (24 years). Exemplar activation maps showing significantly activated voxels (*z* > 3.1) following passive somatosensory stimulation of the right wrist (left). Mean BOLD timeseries (right) extracted from above-threshold voxels, with stimulation periods indicated in grey.

**Table 1.** Neonatal demographics. PMA = post-menstrual age.

|  |  | **Gestational**<br>**age at birth**<br>**(weeks)** | **Post-**<br>**menstrual**<br>**age at scan**<br>**(weeks)** | **Postnatal**<br>**age at scan**<br>**(days)** | **Sex (%**<br>**female)** | **Birthweight**<br>**(kg)** |
| --- | --- | --- | --- | --- | --- | --- |
| **Scanned,**<br>**n = 40** | Median | 35.00 | 37.71 | 15 | 55 | 2.51 |
|  | Range | 28.00–41.57 | 33.57–46.43 | 1–81 |  | 1.17–4.22 |
| **Included,**<br>**n = 23** | Median | 35.00 | 37.00 | 16 | 61 | 2.15 |
|  | Range | 31.29–39.43 | 33.57–45.00 | 1–81 |  | 1.38–4.22 |
| **Preterm,**<br>**n = 8** | Median | 33.43 | 36.00 | 13 | 100 | 1.83 |
|  | Range | 32.14–36.43 | 33.57–36.86 | 3–18 |  | 1.38–2.75 |
| **Early-term,**<br>**n = 8** | Median | 34.43 | 37.43 | 19 | 25 | 2.14 |
|  | Range | 31.29–36.86 | 37.00–38.14 | 4–47 |  | 1.64–2.67 |
| **Late-term,**<br>**n = 7** | Median | 36.29 | 39.71 | 24 | 57 | 2.55 |
|  | Range | 33.43–39.43 | 39.43–45.00 | 1–81 |  | 1.38–4.22 |

Passive right wrist sensorimotor stimulation induced significant clusters of functional activation in the hand area of contralateral (left) S1 in 38/40 neonates and 4/4 adults, consistent with studies at lower magnetic field strengths^1^. The BOLD response timeseries in all significantly activated voxels (*z*-statistic > 3.1) showed a positive change in signal during stimulation periods in all adults and neonates (Figure 1E, F). We excluded 2/40 infants with an absent stimulus specific BOLD response in S1, along with an additional 7/40 neonates due to excessive head motion, 6/40 due to probable stimulus equipment failure, and 2/40 infants with unexpected clinically significant brain pathology, leaving a final dataset of 23 infants. To characterise BOLD response development across the perinatal period, infants were grouped according to PMA (Table 1): preterm (*n* = 8, range = 33.57–36.71 weeks PMA, median = 36.00 weeks PMA); early-term (*n* = 8, range = 37.00–38.14 weeks PMA, median = 37.43 weeks PMA); late-term (*n* = 7, range = 39.43– 45.00 weeks PMA, median = 39.71 weeks PMA).

### Cortical-depth-specific development of the BOLD response to sensorimotor stimulation

In each subject, the hand area of the contralateral (left) S1 was delineated and divided into three cortical-depths (Figure 2). BOLD timeseries across depths were then averaged across each stimulation trial within neonatal and adult groups to derive group trial-average responses (Figure 3). The neonatal BOLD response across all ages was characterised by a delayed rise and return to baseline, in agreement with previous work^9^. Specifically, trial-averaged responses across cortical depths in the preterm group showed a rise in BOLD signal 5 s after stimulus onset, initially peaking around 13 s, with a second, later peak around 27 s before returning to baseline at around 43 s. The BOLD signal amplitude reached a peak of around 3.0%, 1.5% and 1.0% change at superficial, middle and deep cortical depths, respectively. From 37 weeks PMA (early– and late-term groups), the BOLD signal plateaus after a single peak at 13s, before a small post-stimulus undershoot (PSU). The PSU was most pronounced at superficial depths, reaching −0.5% BOLD signal change in the late-term group. In adults, trial-averaged BOLD responses showed a rise around 0–2 s, a much earlier initial peak at 8 s, followed by a plateau and then PSU at 32 s. BOLD signal amplitude reached a peak of around 2.5%, 2% and 1.5% and a PSU minimum of around −0.5%, −0.25% and −0.25% BOLD signal change at superficial, middle and deep cortical depths respectively.

**Figure 2.**
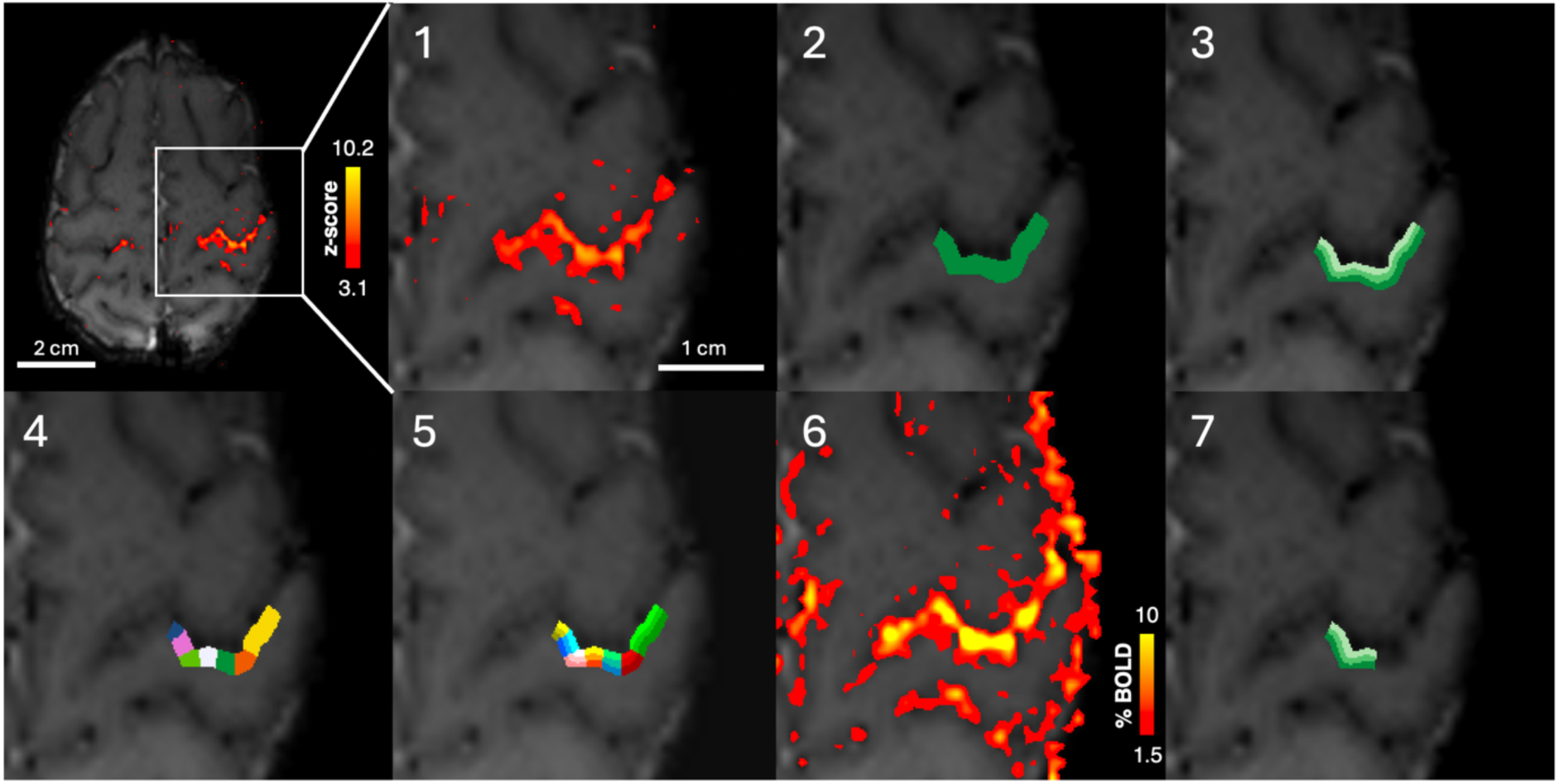
Region of interest (ROI) selection and cortical-depth bin definition for analysis of cortical-depth-dependent BOLD responses. 1. Significantly activated voxels in the contralateral primary somatosensory cortex following somatosensory stimulation of the right wrist. 2. A large ROI is manually defined (green) of the hand area of S1 including the most significantly activated voxels. The ROI spans the full thickness of the cortex across multiple EPI slices. 3. The ROI is split in into three “equivolume layers” or depth bins using LAYNII, where each shade of green indicates a depth bin, with light green being the most superficial depth bin. 4. Each slice is split into seven columns, denoted by colour. 5. The column and layer profiles are combined to produce columns with three equivolume cortical-depth bins, with colours corresponding to columns and shades corresponding to depth bins. 6. The activation map in units of BOLD percent signal change. 7. The columns in which the BOLD percent signal change in the deep cortical-depth bins was in the top 40^th^ centile were selected to produce the final ROI used for the cortical-depth-dependent analyses, with depth bins denoted by shades, with the lightest shade representing the most superficial depth bin.

**Figure 3.**
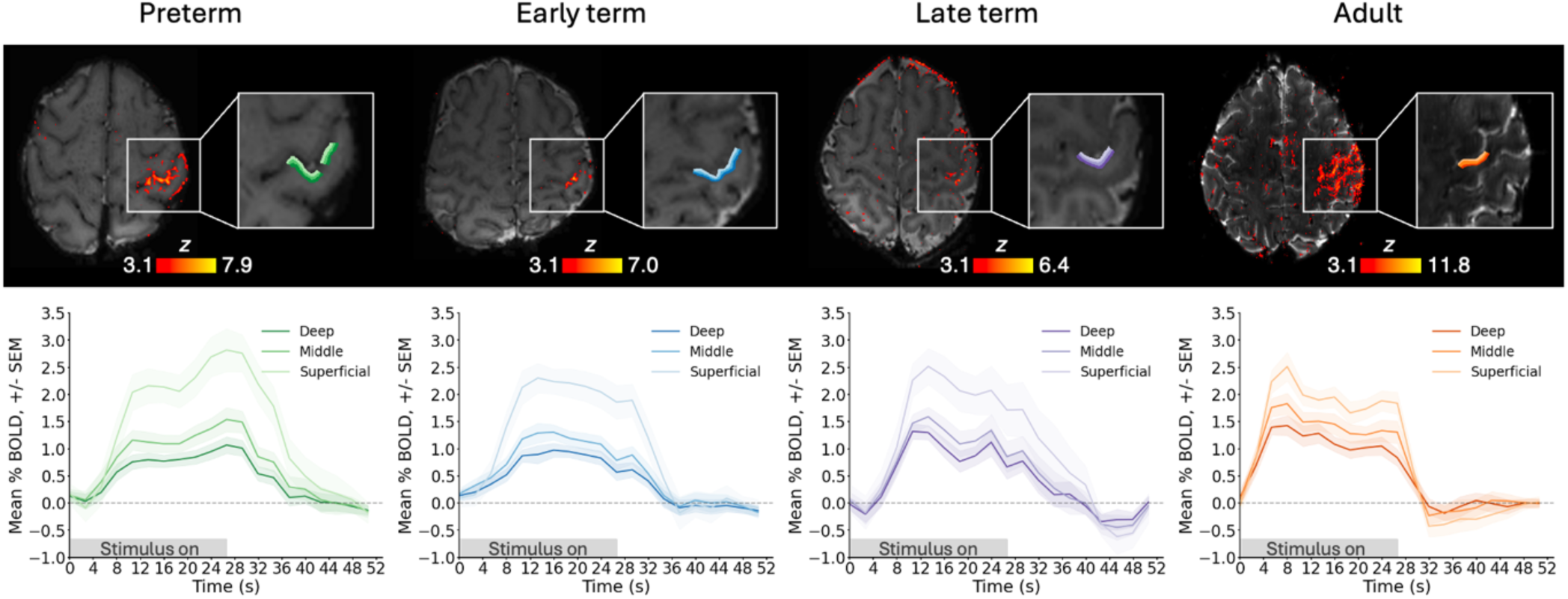
The cortical-depth-dependent BOLD response in S1 to passive somatosensory wrist stimulation is immature in the neonatal period and develops with age. (Top) Exemplar significantly activated voxels (*z*>3.1) following sensorimotor stimulation to the right wrist and three equivolume cortical depths in the hand area of contralateral primary somatosensory cortex (S1) overlaid onto the mean EPI data. Lightest shade denotes the most superficial depth. Trial-average cortical depth dependent BOLD response timeseries (+/− SEM) in preterm, (*n*=8 neonates), early-term (*n*=8), and late-term (*n*=7) neonates, and adults (*n*=4).

A positive BOLD signal change was seen at all cortical depths in contralateral S1 across every trial in both neonates and adults, irrespective of age. BOLD activity was predominantly maximal at the most superficial depth and decreased towards the deeper cortical grey-white matter boundary (Figure 4). The ratio of maximum BOLD signal change across cortical depths in each trial (Figure 3B) was significantly different across age groups in the superficial:middle (*H*(3) = 35.12, *p* <0.001) and superficial:deep ratios (*H*(3) = 35.96, *p* <0.001), but not for the middle:deep ratio (*H*(3) = 7.06, *p* = 0.070). A post-hoc Dunn test was significant for decreasing superficial:middle between all neonatal age groups and adults (preterm vs. adults: *p* <0.001, early-term vs. adults: *p* <0.001, late-term vs. adult: *p* = 0.024), and approaching significance for early-term vs. late-term (*p* = 0.091). Superficial:deep significantly decreased between preterm vs. adults (*p* <0.001), early-term vs. late-term (*p* = 0.041), early-term vs. adults (*p* <0.001), and approaching significance between late-term vs. adults (*p* = 0.104) (for full results see Tables S1–3). This indicates that the change in stimulus-associated BOLD response develops in a cortical-depth-specific manner.

**Figure 4.**
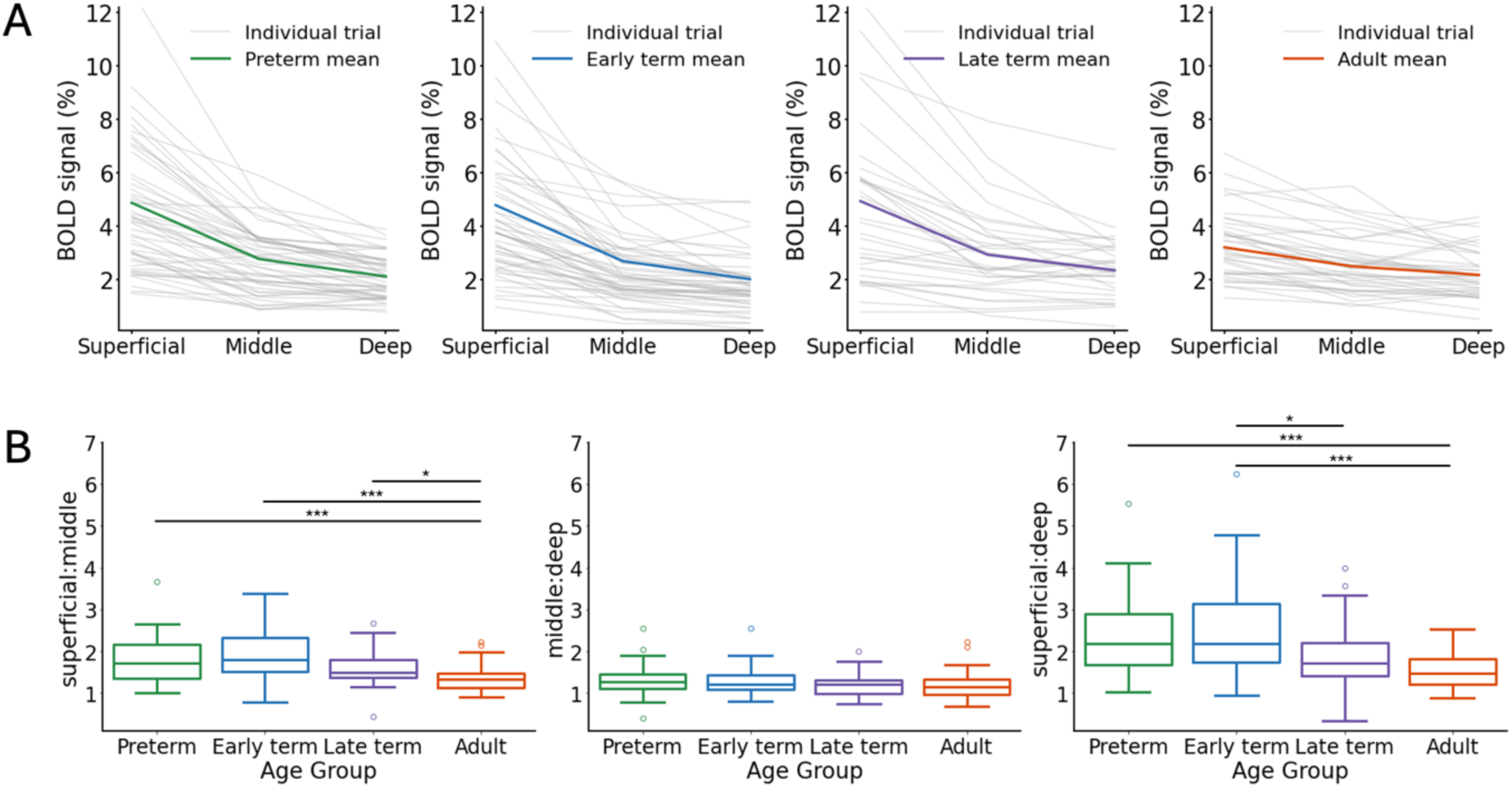
Maturation of the BOLD response is cortical-depth specific. (A) The cortical-depth profile of the maximum BOLD percent signal change across cortical depths averaged across all trials in preterm (*n*=8), early-term (*n*=8), and late-term (*n*=7) neonates, and adults (*n*=4). (B) Significantly dijerent ratios of maximum BOLD precent signal change were found across age groups for Superficial:Middle (Kruskal-Wallis: H(3) = 35.12, *p* <0.001. Dunn post-hoc with Bonferroni adjustment: preterm vs. adult: *p* <0.001, early-term vs. adult: *p* <0.001, late-term vs. adult: *p* = 0.024) and superficial:Deep (Kruskal-Wallis: H(3) = 35.96, *p* < 0.001. Dunn post-hoc with Bonferroni adjustment: preterm vs. adult: *p* <0.001, early-term vs. late-term: *p* = 0.041). (* indicates *p* <0.05; ** indicates *p* <0.01; *** indicates *p* 0.001.)

To investigate whether the increased ratio of superficial to deeper depth BOLD signal change in neonates could simply be explained by a different extravascular BOLD signal extent around large pial surface veins, we applied the infinite-cylinder model of extravascular field perturbations^21,22^. As neonatal cortical tissue is more water-rich^23^ and has a lower iron concentration^24^ compared with adult tissue, neonatal grey matter tissue susceptibility should be markedly reduced compared to adults. Thus, blood-tissue susceptibility differences (Δχ_blood-tissue_) would be larger, in turn altering extravascular field properties. Accounting for the different neonatal tissue properties by increasing the adult Δχ_blood-tissue_ by a factor of 2, 3 and 4 (0.5, 1, 1.5 and 2 ppm respectively; Table 3) resulted in simulated field perturbations that were negligible (0.02 ppm) at a distance of 0.08, 0.13, 0.17 and 0.2 mm respectively from the vessel (Figure 5A). A further consideration is that neonatal large pial veins are half the diameter of adult veins^25^. Field perturbation simulations using adult-like and neonate-like large pial vein diameters and Δχ_blood-tissue_ (Table 4) demonstrated that magnetic field perturbations reach negligible levels (0.02 ppm) around the adult-like vessel at a distance of 0.2 mm, which was reduced to 0.13 mm for the neonate-like vessel (Figure 5B). Together this suggests that despite developmental differences in vascular architecture, tissue and blood properties, the spatial extent of extravascular fields from large pial veins are smaller in neonates compared with adults. The proportionally increased BOLD amplitude at superficial depths cannot therefore be attributed to the spatial extent of extravascular fields, and instead likely reflects age-specific densities of blood vessels across different cortical depths.

**Figure 5.**
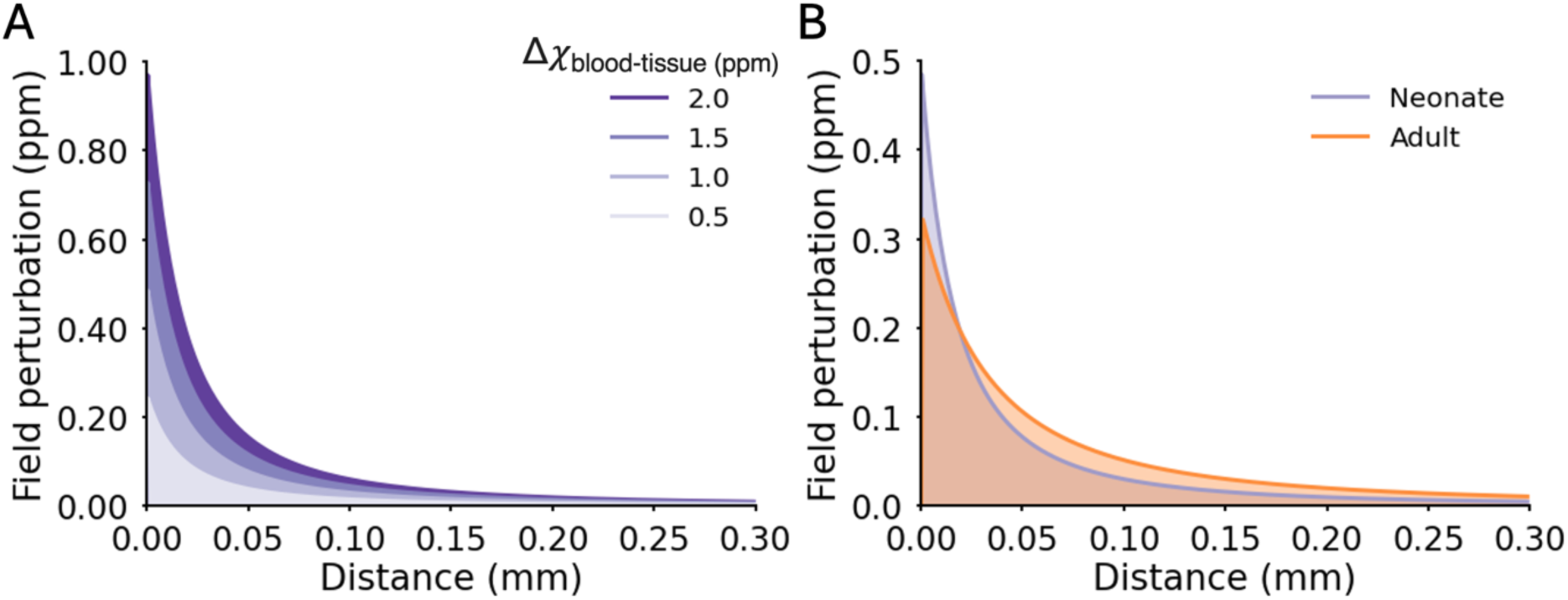
Spatial extent of extravascular fields of large pial veins is not larger in the neonatal period compared with adults. (A) The simulated field perturbation produced for a 65-µm diameter pial vein oriented perpendicular to B_0_ using the infinite-cylinder model of extravascular fields. A range of blood-tissue susceptibility (Δχ_blood-tissue_) values (0.5, 1.0, 1.5 and 2.0 ppm) were simulated with 0.5 ppm being the adult-like value. A distance of 0 mm denotes the boundary between the vein and the grey matter tissue. (B) The predicted field perturbation for an adult-like large peripheral vein (orange; 65-µm diameter and 0.5 ppm Δχ_blood-tissue_)and a neonate-like large peripheral vein (purple; 130-µm diameter and 1 ppm Δχ_blood-tissue_).

**Table 2.** Functional MRI sequence parameter values for data acquisition in neonatal and adult participants.

| Parameter | Adult | Neonate |
| --- | --- | --- |
| Acquisition | 2D GRE-EPI BOLD | 2D GRE-EPI BOLD |
| Resolution | 0.8 mm isotropic | 0.8 mm isotropic |
| Matrix size | 260 x 260 x 57 | 156 x 156 x 25 |
| Fourier | 5/8 | Full |
| Repetition time (TR; ms) | 2660 | 2660 |
| Echo time (TE; ms) | 30 | 48 |
| Flip angle (°) | 45 | 90 |
| Nominal echo spacing (ms) | 1 | 1.06 |
| Inplane-acceleration factor | 2 | 2 |
| Slice-acceleration factor | 3 | N/A |
| Acquisition time | 11 m 5 s | 11 m 5 s |

**Table 3.** Infinite-cylinder model parameter values extravascular field simulations investigating the eject of increasing susceptibility dijerence between cortical tissue and venous blood (Δχ_blood-tissue_).

| Parameter |  |
| --- | --- |
| $\Delta\chi_{\text{(blood-tissue)}} \text{ (ppm)}$ | 0.5, 1.0, 1.5, 2.0 |
| Vessel radius ($\mu\text{m}$) | 32.5 |
| Angle of vessel to $B_0$, $\theta$ | 90 |
| Magnetic field strength, $B_0$ (T) | 7 |

**Table 4.** Infinite-cylinder model parameter values for adult-like and neonate-like extravascular field simulations.

| Parameter | Adult | Neonate |
| --- | --- | --- |
| Vessel radius ($\mu\text{m}$) | 65 | 32.5 |
| $\Delta\chi_{\text{(blood-tissue)}} \text{ (ppm)}$ | 0.5 | 1.0 |
| Angle of vessel to $B_0$, $\theta$ | 90 | 90 |
| Magnetic field strength, $B_0$ (T) | 7 | 7 |

## Discussion

We present the first evidence of depth specific haemodynamic responses in the human neonatal cerebral cortex. Maturational differences were seen both across the neonatal period and when compared to adults. Passive right wrist sensorimotor stimulation induced spatially-specific positive BOLD responses in the cortical grey matter, indicative of functional hyperaemia in the neonatal brain from as early as 33 weeks PMA. Consistent with prior findings, we found that BOLD response temporal features were also altered during the neonatal period^9^. We further demonstrate that the increased spatial resolution and fCNR afforded by fMRI at 7 T enables discrimination of BOLD responses across cortical depths in the neonatal period, despite reduced cortical thickness^20^. Significant age-specific differences in the ratios of maximum BOLD signal change across cortical depths suggest that BOLD response development is not simply due to global neonatal physiology changes. Instead, multiple mechanisms underpinning the BOLD response — such as neurovascular coupling, vascular anatomy and neuronal activity itself—are all likely evolving in a depth-specific manner within the developing cortex prior to and around the time of full-term birth.

The time to onset and peak of the neonatal BOLD response was delayed in all neonatal groups compared to adults, as seen in previous neonatal human^9^ and juvenile rodent studies^26^. This is consistent with evidence showing neuronal responses to somatosensory stimulation undergoing rapid decreases in latency and duration across the last trimester of gestation and early infancy^27,28^. Given the markedly different timescales of developmental changes in neuronal responses (in the order of 100 ms) and the BOLD response (in the order of seconds), the delayed time to peak observed here in neonatal BOLD responses is unlikely to be due to differences in neuronal processing alone, and likely relates to concomitant immature haemodynamics and/or neurovascular coupling.

Onset timing of the BOLD response is largely dictated by the speed of oxygenated blood delivery, which is determined both by baseline cerebral blood flow (CBF) and cerebrovascular reactivity. Resting neonatal CBF is around one third of that in adults^9,29^. This is likely due to cortical vascular development, as diving arteriole diameter increases across the mouse equivalent of the human third trimester produces increases in resting CBF^30^, while arterial muscularisation also increases capacity for dynamic CBF regulation and propagation of dilation upstream from the capillaries^31^. Moreover, other non-neuronal components of the complex neurovascular coupling cascade are present but also relatively immature^2^. For example, in mice, although astrocytes regulate cerebral vessel diameter as early as the equivalent of full term in human neonates^32^, their number, size, maturity and associated synapses are all increasing up to birth, with mature ramified astrocyte morphology not emerging until well after the neonatal period^33^. Maturation of cortical vasculature and non-neuronal components of the neurovascular coupling cascade thus likely act together to reduce BOLD response onset times across development.

In data collected from our youngest infants, BOLD responses exhibited a second, longer-latency positive peak after the first, initial positive peak. A second longer-latency peak is also seen in electrophysiological visually-evoked responses in preterm infants and juvenile rodents which disappears as development proceeds^34^. We similarly see a transition in the early-term period, after which the BOLD response exhibits a more canonical decrease and plateau after the initial, early positive peak, as in adults. The plateau is thought to reflect adaption of neuronal activity to a repeating stimulus, mediated by inhibitory circuits^35,36^. In neonates, integration of inhibitory GABAergic neurons into cortical circuits is ongoing and thus interneuron firing patterns are still immature^37^. Thus, inhibitory neurons may not influence cortical neuronal activity to the same extent in the younger infants, resulting in an absent BOLD response plateau. Non-neuronal mechanisms (particularly hemodynamic) could also explain the biphasic BOLD response of the youngest infants; as in rodents there is a late “feedback” component of neurovascular coupling multiple seconds after the onset of functional hyperaemia^38^, and a glial calcium transient occurring several seconds after the initial fast neuronal calcium transient^39^.

Following the stimulus-induced BOLD response positive peak, we observed a small negative dip starting in the late-term infants, reminiscent of the BOLD PSU commonly observed in adults. The BOLD PSU reflects a later transient local increase in deoxygenated haemoglobin concentration, although the exact biological mechanism remains debated^40^. One proposed mechanisms is venous ballooning due to higher vessel compliance^41^. In neonates, vessel wall composition is still maturing^42^ and mechanical properties of the parenchymal tissue are altered due to more water-rich and less cell-dense tissue composition^23^. Thus, mechanical constraints on vascular dilation and vessel ballooning are likely altered and may affect the PSU^43^. Alternatively, prolonged elevation in CMRO_2_ persisting after CBF and CBV have returned to baseline following neuronal activation have also been proposed to underlie the BOLD PSU^44^. This is relevant to our study period, as there is rapid, wide-scale cortical synaptogenesis across development leading to maturational changes in baseline and activity induced increases in cortical metabolism and CMRO_2_^45,46^. Elevation in CMRO_2_ following stimulus cessation may thus increase leading to the emergence of a BOLD PSU in infants.

The BOLD signal measured with fMRI is composed of intravascular and extravascular contributions, however at 7 T, the signal measured with standard GRE methods is largely due to extravascular BOLD^47^. Local regulation of blood flow, at the level of the parenchymal microvasculature, makes fine-scale mapping of neuronal activity across the cortex possible. A key consideration of GRE-BOLD is that spatial specificity can be hampered by large extravascular fields around draining pial veins distal to the neuronal activity^10^. The proportionally larger superficial depth BOLD response amplitude in neonates could suggest that a larger pial-vein bias is present in neonatal GRE-BOLD fMRI data at 7 T. However, despite larger susceptibility differences between blood and neonatal cortical tissue^24^, we found the spatial extent of extravascular fields in the neonatal cortex is reduced compared with that in mature adult cortex. This is because smaller pial vein diameters in neonates offset effects due to larger blood-tissue susceptibility differences compared to adults. Therefore, with cortical thickness estimated to be around 1.5–2 mm and field perturbations due to extravascular fields negligible by a distance or depth of 0.13 mm, we expect minimal coupling of the BOLD responses measured at the pial surface and deeper within the parenchyma in neonates.

The neonatal pial vasculature comprises a dense anastomotic plexus of capillary-like vessels covering the pial surface, which undergo rarefaction postnatally^48^. The increased vessel density on the pial surface may therefore produce a larger amplitude BOLD response on the pial surface during this period as seen in our data. Intense angiogenesis occurs in the cortical grey matter, firstly at the deeper depths then progressing towards the surface^49^, to rapidly generate the dense intraparenchymal capillary network seen in the adult brain^50^. In the period equivalent to the last trimester of gestation, this largely comprises capillary angiogenesis and higher-order capillary sprouting, more than doubling parenchymal vessel density in mice^48^. Importantly, despite the presence of an adult-like blood vessel hierarchy in the cortex around the time of birth, blood flow regulation across this hierarchy is likely still developing, including pericyte integration and maturation at the level of capillaries^51^ and arterial smooth muscle^31^. Redistribution of relative vessel density and therefore cerebral blood volume across the depths of the cortex, alongside maturation of blood flow regulatory mechanisms across the vascular architecture may increase BOLD response amplitude in middle and deeper regions while decreasing superficial amplitudes across development.

Decreased levels of neuronal activity and immature cortical circuitry may also contribute to comparatively smaller BOLD signal amplitudes at the deep and middle cortical depths in early development. Widespread synaptogenesis begins around mid-gestation^52^ and continues until early childhood^53^, as reflected by increasing local field potential (LFP) amplitudes in mice following sensory stimulation^26^. As a result, baseline metabolism and CMRO_2_ increase across this same period of development^45,54^. This suggests that the immature neuronal response to sensory stimulation is likely diminished in neonates compared with adults, resulting in smaller amplitude BOLD responses specifically in the deep and middle cortical depths.

BOLD signal specificity, sensitivity and amplitude may also be modified because of how immature cerebral physiology interacts with the physics of the BOLD signal. Our results show that distinct activation amplitudes can be resolved across cortical depths, such that the pial surface signal can be isolated and separated from the parenchyma. Consistencies in temporal features of the BOLD response across cortical depths suggests that we may not be specifically sampling from distinct levels of the vascular hierarchy at each depth, and therefore it is unclear whether responses are depth-specific enough to infer cortical ‘functional circuitry’. Alternatively, the temporal resolution used here may be too coarse to identify subtle differences in temporal features across depths. Tissue and blood composition are also different in the neonatal brain, with increased water and decreased tissue cell density, immature myeloarchitecture^23^ and elevated blood haematocrit and haemoglobin^55^. Together this presents an implicit confound due to the co-development of brain activity, cerebral physiology and their interaction with MRI physics, representing a clear challenge for specific interpretation of BOLD fMRI signal changes across age.

Consistent with most neonatal task-based fMRI studies, data were acquired from infants during natural sleep to minimise head motion whereas adults were scanned awake. Sleep is the main behavioural state in newborn infants (representing up to 80% of their day), encompassing two main sleep states^56^: quiet sleep and active sleep. Although neuronal responses to visual stimulation during sleep and wake states are identical in preterm infants^34^, sleep has been found to cause large haemodynamic fluctuations that confound fMRI responses in neonatal mice, particularly during sleep-state transitions^57^. As stringent motion exclusion criteria (<0.8 mm) were used, it is likely that the neonates studied here remained in quiet sleep (precursor to NREM sleep) for the duration of the trials included, however simultaneous EEG recordings would be required to definitively characterise how infant sleep states may affect the measured stimulus-evoked BOLD responses.

Due to higher predicted specific absorption rate (SAR) levels in neonates compared with adults under equivalent scanning conditions at 7 T^18^, more conservative approaches were required for the fMRI acquisition, including reduced SAR intensive acquisitions. Scan duration is largely limited in infants to how long they remain in natural sleep, which is typically around 50 minutes^17^. Irrespective of sleep, following local safety assessments, scan sessions were limited to 1 hour in infants to avoid possible temperature instability—both hypothermia due to reduced subcutaneous fat and temperature regulation capabilities in early life, as well as systemic heating due to SAR^17^. As a result, the number of functional runs acquired were limited compared with adult 7 T fMRI studies, which regularly use multiple functional runs or even scanning sessions. Furthermore, unconstrained gross head motion is a more significant challenge when scanning infants compared with adults, leading to more data being discarded due to motion corruption. Differences in tissue properties of the neonatal brain, as well as relative size of brain structures and SNR, meant that careful optimisation of preprocessing and analysis approaches was required to ensure the tools were working appropriately for 7 T neonatal fMRI data. Here, a minimal data processing approach was used to conserve spatial specificity of the signal. As more neonatal 7 T fMRI experiments are carried out, further work would be required to further optimise the full complement of tools regularly used for different fMRI preprocessing and analysis approaches to account for the unique properties of neonatal 7 T fMRI data.

The ability to identify cortical-depth-dependent differences in the BOLD response in neonates has significant implications for the study of cortical circuit formation during the crucial period around normal birth. This can include characterisation of how processing of distinct stimuli occurs across circuitry localised to the different anatomical layers of the cortex. In the mature brain this has provided insight into the processing of different visual stimuli across depths^58^ and differences in input– and output-driven activity in the primary motor cortex (M1) during different finger tapping tasks^13^. Laminar functional connectivity profiles can also be defined at rest, demonstrating stronger functional connectivity between the deeper laminae of M1 with the premotor area and between the superficial laminae of M1 and S1^13^. In the last trimester of human gestation, the large-scale cortical connectivity underpinning sensory processing is still emerging^1^, with evidence of hierarchical processing first seen from 34 weeks PMA with ongoing development beyond term equivalent age^59^. Characterisation of activation and functional connectivity across cortical depths will thus enable a better understanding of how functional specialisation and its associated circuitry across anatomical cortical layers first emerges across development in sensory cortices. This is critical as cortical circuitry is often disrupted in neurodevelopmental conditions, with altered sensory processing being a key characteristic of many such conditions^60^.

Over the past 20 years, there has been ongoing uncertainty regarding whether functional hyperaemia is even present during development, with BOLD fMRI, functional near-infrared spectroscopy, and invasive optical imaging methods in neonatal humans^9,61^ and juvenile rodents showing the presence of positive, negative and absent BOLD responses^26,62^. Altered functional hyperaemia in early development has been proposed to relate to a variety of mechanisms including: vasoconstriction of pial arteries^62^; systemic blood pressure changes and a lack of cerebrovascular homeostasis resulting in positive BOLD responses that do not reflect local neuronal activity^62^; or decreases in regional blood flow^63^. Here, we find local, spatially specific positive BOLD responses from as early as 33 weeks PMA in response to a somatosensory stimulus. This is indicative of spatially tuned functional hyperaemia in the neonatal human brain, which would not be consistent with the aforementioned absent of negative BOLD responses in development. Similar to other previous development studies^9,26,62,63^, we demonstrate an increase in positive BOLD response amplitude and a decrease in the onset latency of the BOLD response with increasing age. To understand this process further, it would be insightful to perform studies with even younger neonates, to determine when and how spatially specific functional hyperaemia first emerges.

## Conclusion

Here we demonstrate the feasibility of characterising functional activation across cortical depths in the neonatal brain using ultra-high-field BOLD fMRI. We demonstrate that the BOLD response in neonates across the cortical vascular hierarchy differs significantly from the adult response. Accounting for known anatomical vascular differences between neonates and adults suggests that our observations are likely predominately driven by physiological changes during development. Precise characterisation of the neonatal BOLD response with such high spatial specificity thus provides a unique opportunity to establish fundamental insights into how brain activity and its associated framework of circuitry is established in the human cortex during this critical period of early brain development. As a result, it may lead to a greater understanding of how disruption to this critical process results in neurodevelopmental disorders or neurological disease manifesting in adulthood.

## Methods

### Participants

Data were collected from 40 neonates aged 33.57–46.43 weeks PMA (median: 37.71 weeks PMA, 22 female) recruited from the postnatal and neonatal wards at St. Thomas Hospital, London (Table 1). Data were also collected from 4 healthy adult volunteers aged 22–27 years old (median: 25.5 years, 3 female). UK National Health Service ethics committee approval (NHS REC: 19/LO/1384) and written consent (from parents in the case of neonatal participants) was acquired prior to data collection.

### Subject preparation

Neonates were fed and then swaddled in two pre-warmed blankets and immobilized using a vacuum evacuated blanket (Pearltec, Zurich, CH), with the head positioned in the isocentre of the head coil with the aid of foam cushions^17^ (Figure 1A). To provide hearing protection, moulded dental putty (President Putty, Coltene Whaledent, Mahwah, NJ, USA) was fitted in the external auditory meatus of the infant’s ear and inflatable cushions were placed over the ears which also minimized head motion (Pearltec). Neonates were scanned for a maximum of 1 hour following feeding and during natural sleep, with concurrent temperature and vital sign monitoring carried out by an experienced clinical staff member.

Adults were positioned supine headfirst in the head coil. Foam ear plugs were used to provide hearing protection and were scanned awake at rest for a maximum of 1 hour.

### MRI data acquisition

Images were acquired on a Siemens 7 T system (MAGNETOM Terra, Siemens Healthineers, Erlangen, DE) with a 1Tx-32Rx and 8Tx-32Rx Nova Medical head coil array (Wilmington, MA, USA) in neonates and adults respectively. We used an in-house image-based B_0_ shimming method, requiring a low-resolution whole-head B_0_ field map, which was then fitted with empirically-calibrated basis functions, similar to the approach discussed by D’Astous et al^64^. This approach was found to significantly outperform the built-in vendor-provided solution.

fMRI data were acquired using a single-shot gradient-echo EPI sequence with Dual-Polarity GRAPPA (DPG) reconstruction^65^ over a restricted field-of-view including M1 and S1 in both hemispheres (Figure 1D). Acquisition parameter values were kept as consistent as possible between adult and neonatal acquisitions, but there were some differences, primarily to account for the difference in brain size and brain T_2_* of neonates compared with adults^66^ (see Table 2 for details of protocol parameter values). Recent detailed RF safety simulation demonstrated higher SAR levels in neonates at 7 T compared with adults under equivalent scanning conditions^18^. In accordance with local risk assessments, the scanner software was modified to use more conservative SAR estimation when imaging neonates^17^.

High-resolution 2D T_2_-weighted (T_2_w) images were also acquired in at least two orthogonal planes using a turbo spin echo (TSE) sequence with parameters 0.6 mm × 0.6 mm × 1.2 mm, TR/TE 8,640/156 ms and an inplane-acceleration factor of 2. Orthogonal T_2_w images were motion corrected and reconstructed to an isotropic resolution of 0.45 mm using SVRTK (https://github.com/SVRTK/SVRTK)^67^. All high-resolution structural images were formally reviewed and reported by an experienced Neonatal Neuroradiologist to identify any pathology and check for normal appearance for age.

### Sensorimotor stimulus

To provide safe and reproducible sensorimotor stimulation to the right wrist inside the MRI scanner, infants were fitted with a custom-built 3D-printed MR-compatible robotic device specifically sized to the wrist of neonates^68^ (Figure 1B). A pneumatic piston, driven using the hospital-pressurised air supply, moved the robotic device and therefore the wrist through a smooth continuous pattern of flexion and extension at a frequency of 0.5 Hz, time-locked to the fMRI acquisition by receiving the volume TTL trigger pulse using a data acquisition unit (Labview, National Instruments, Austin, TX, USA). Each experiment involved an “on-ob” boxcar paradigm consisting of 7–12 blocks of 26.6-s of wrist stimulation interleaved with 26.6-s of rest. The same device that was used in neonates was also used in adults, after being adapted using LEGO (Billund, DK) to better fit the adult hand (Figure 1C).

### fMRI data preprocessing and analysis

Neonatal and adult fMRI data were preprocessed using the same pipeline. Firstly, fMRI data were corrected for differences in acquisition time between slices within the same volume using the slicetimer from the FSL software package (https://www.fmrib.ox.ac.uk/fsl)^69^, interpolating the data to the middle slice of each volume. Excessive motion was detected using MCFLIRT from FSL^70^, and data were cropped to exclude corrupted volumes if motion occurred at beginning or end of the acquisition. Thermal noise components were removed from the data using NOise Reduction with DIstribution Corrected (NORDIC) principle component analysis (v1.1; https://github.com/SteenMoeller/NORDIC_Raw)^71^ on the magnitude data, and data were motion corrected by registering all volumes to the middle frame using 3dVolReg from the AFNI software package (https://afni.nimh.nih.gov/)^72^ via the mc-afni3 wrapper provided by FreeSurfer (https://surfer.nmr.mgh.harvard.edu/)^73^. If the middle volume was motion corrupted, a volume from an adjacent motion-free period was selected as the alignment target. To further reduce the effect of motion in the data, motion parameters (roll, pitch and yaw in degrees, and column, row and slice displacement in mm) from the motion correction step were regressed out of the fMRI data using the regfilt tool from FSL^69^. The data were also high-pass filtered at 0.02 Hz to remove low-frequency signal drifts.

Activation associated with the wrist stimulus was identified using a general linear model (GLM; implemented in FSL FEAT) timeseries analysis with pre-whitening (FSL FILM) and the stimulus timing convolved with neonatal-^9^ or adult-specific haemodynamic response basis functions^74^. An additional confound explanatory variable of volumes with excessive motion was also used, as identified by the root-mean-square (RMS) intensity difference compared with the reference (first) volume using fsl_motion_outliers from FSL^69^. The GLM produced a *z*-statistic activation map thresholded at *z* > 3.1. The parameter estimates of the BOLD response from the GLM were then converted to BOLD percent signal change (https://jeanettemumford.org/assets/files/perchange_guide.pdf).

### Cortical depth-dependent analysis

A region of interest (ROI) was manually defined on the mean functional image in the hand area of contralateral S1 encompassing the cortical grey matter voxels with the largest *z*-scores (*z* > 3.1) associated with the wrist stimulus and spanning the whole thickness of the cortex. EPI data were then resampled using 3dResample from AFNI^72^ to a higher spatial resolution (0.16 mm × 0.16 mm × 0.8 mm). The ROI was refined at this higher resolution to ensure the pial edge and grey/white matter boundary had been accurately delineated. The refined ROI was then segmented into three “equivolume layers” or cortical depth bins and each slice of the ROI segmented into seven columns, with both layers and columns defined using LAYNII^75^. To circumvent the inherent pial bias of the BOLD signal in our final ROI selection, BOLD percent signal change was extracted from the deepest depth in each column, and columns in the top 40^th^ centile of percent signal change at this depth were selected for the final ROI (Figure 2).

To investigate how the BOLD response timeseries developed with PMA, the neonatal data were split into three groups based on age: preterm (*n* = 8, range = 33.57– 36.71 weeks PMA, median = 36.00 weeks PMA), early-term (*n* = 8, range = 37.00–38.14 weeks PMA, median = 37.43 weeks PMA), late-term (*n* = 7, range = 39.43–45.00 weeks PMA, median = 39.71 weeks PMA). The whole timeseries was extracted from each depth bin in the final ROI and the data were converted to BOLD percent signal change using the first stimulus-on TR and the last stimulus-ob TR from that stimulus trial (10 TRs stimulus on, 10 TRs stimulus ob) in neonates and using the last two stimulus-ob TRs in adults as the baseline. The mean BOLD percent signal change timeseries was calculated in each cortical depth bin from each subject in each age group to give a group-level trial-averaged BOLD response to the sensorimotor wrist stimulus. Trials were excluded from the trial-averaged response if displacement with respect to the reference volume was greater than 0.8 mm (one voxel).

### Statistical analyses

The maximum BOLD percent signal change in each cortical depth bin in each trial was taken from each age group and the ratio of maximum BOLD percent signal change was calculated between each cortical depth bin. Although the maximum signal change can be slightly unstable due to random variance introduced by noise, we opted to use the maximum because it is model-free, avoiding assumptions in response timing, compared with a more complex model using a range of “active” values. Furthermore, the maximum BOLD percent signal occurred at similar relative timepoints across each cortical depth bin. A Kruskal-Wallis test with a Dunn post-hoc test and Bonferroni adjustment for multiple comparisons was used to test for a significant difference in cortical depth bin ratios between age groups.

### Modelling extravascular fields

To predict the extravascular field perturbations produced by the susceptibility difference of venous blood and the surrounding tissue, we modelled a single blood vessel as an infinite-cylinder which produces magnetic field inhomogeneities in the surrounding grey matter tissue^21,22^. The characteristics of these extravascular fields are dependent on properties of the blood, the surrounding tissue, the orientation of the vessel to the main magnetic field and the vessel itself. The field offset is given by

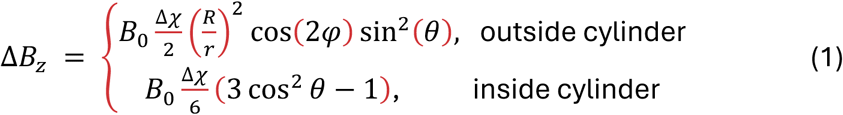

where the difference in susceptibility difference (Δχ) is the blood-tissue susceptibility difference given by

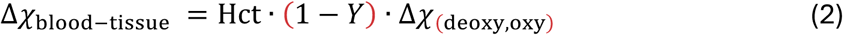

and is dependent on the blood haematocrit (Hct) and blood oxygenation (*Y*). The susceptibility difference between fully-oxygenated and fully-deoxygenated blood (Δχ_(deoxy,oxy)_) has been previously reported as 0.264 ppm^76^.

The magnetic field inhomogeneities produced inside and outside of the a blood vessel (cylinder) can be described by equation (1) where Δ*B*_*z*_ is the perturbation of the main magnetic field (*B*_”_) produced at a point in space described by the vector *r* perpendicular to the vessel with a magnitude *r* for a vessel of radius (*R*) oriented at an angle (θ) to *B*_”_ with a given susceptibility difference (Δχ). Here, Δχ is the difference in susceptibility between the whole blood in the vessel and the surrounding tissue (Δχ_blood-tissue_) is described in equation (2).

The infinite-cylinder model was simulated for single blood vessel in a field of strength 7 T to give the field inhomogeneities perpendicular to the blood vessel with increasing distance from the vessel wall. A vessel perpendicular to *B*_”_was used to produce the maximal extravascular field, assuming all other parameters held constant^21,77^. As cortical tissue is much more water-rich, less cell-dense and contains less iron^23^ in the neonatal period, it would be expected to have a greater Δχ_blood-tissue_ compared with venous blood than in adults. To quantify the effect of this as well as reduced pial vein dimeter in neonates, the infinite-cylinder model was adapted with age-specific literature values for (Δχ_blood-tissue_ (Table 3) and large pial vein radius^25,78^ (Table 4).

## Extended Data

**Table S1.** Kruskal-Wallis test to identify age-related dijerences in the ratio of maximum BOLD signal change across cortical depths in each trial between groups.

| Ratio | Kruskal-Wallis test |  |
| --- | --- | --- |
|  | *H*-statistic, *H*(3) | *p*-value |
| Superficial:Middle | 35.12 | <0.001 |
| Middle:Deep | 7.06 | 0.070 |
| Superficial:Deep | 35.96 | <0.001 |

**Table S2.** Post-hoc Dunn test (with Bonferroni correction for multiple comparison) to identify age-related dijerences in ratio of maximum BOLD signal change between superficial and middle depths.

| Superficial:Middle (*p*-values) |  |  |  |  |
| --- | --- | --- | --- | --- |
|  | Preterm | Early-term | Late-term | Adult |
| Preterm | 1.000 |  |  |  |
| Early-term | 1.000 | 1.000 |  |  |
| Late-term | 1.000 | 0.091 | 1.000 |  |
| Adult | <0.001 | <0.001 | 0.024 | 1.000 |

**Table S3.** Post-hoc Dunn test (with Bonferroni correction for multiple comparison) to identify age-related dijerences in ratio of maximum BOLD signal change between superficial and deep depths.

| Superficial:Deep (*p*-values) |  |  |  |  |
| --- | --- | --- | --- | --- |
|  | Preterm | Early-term | Late-term | Adult |
| Preterm | 1.000 |  |  |  |
| Early-term | 1.000 | 1.000 |  |  |
| Late-term | 0.174 | 0.041 | 1.000 |  |
| Adult | <0.001 | <0.001 | 0.104 | 1.000 |

## Acknowledgements

We thank the patients and families who agreed to participate in this work, and the staff of St. Thomas’ Hospital London. J.W.M was supported by PhD funding from the UK Medical Research Council [MR/P502108/1]. T.A. was supported by an MRC Clinician Scientist Fellowship [MR/P008712/1] and a Transition Support Award [MR/V036874/1]. J.W.M and T.A. were additionally supported by an MRC Senior Clinical Fellowship [MR/Y009665/1]. The work was also supported by a project grant awarded by Action Medical Research [GN2728]. J.W.M., P.C., A.D.E., and T.A. received support from the Medical Research Council Centre for Neurodevelopmental Disorders, King’s College London [MR/N026063/1]. G.H. and J.R.P. received support from the National Institutes of Health and the National Institute of Biomedical Imaging and Bioengineering [P41-EB030006 and R01-EB032746]. G.H. also received support from the BRAIN initiative [F32-MH125599, R01-MH111419, U19-NS123717] and the German Deutsche Forschungsgemeinschaft [DFG Project number 543670971]. The authors acknowledge support from a Wellcome Trust Collaboration in Science Award [WT 201526/Z/16/Z], the Wellcome/Engineering and Physical Sciences Research Council Center for Medical Engineering at King’s College London [WT203148/Z/16/Z], and by the National Institute for Health Research (NIHR) Clinical Research Facility based at Guy’s and St. Thomas’ NHS Foundation Trust and King’s College London. The views expressed are those of the author(s) and not necessarily those of the NHS, the NIHR, nor the Department of Health and Social Care.

